# Robust and Stable Gene Selection via Maximum-Minimum Correntropy Criterion

**DOI:** 10.1101/029538

**Authors:** Majid Mohammadi, Hossein Sharifi Noghabi, Ghosheh Abed Hodtani, Habib Rajabi Mashhadi

## Abstract

One of the central challenges in cancer research is identifying significant genes among thousands of others on a microarray. Since preventing outbreak and progression of cancer is the ultimate goal in bioinformatics and computational biology, detection of genes that are most involved is vital and crucial. In this article, we propose a Maximum-Minimum Correntropy Criterion (MMCC) approach for selection of biologically meaningful genes from microarray data sets which is stable, fast and robust against diverse noise and outliers and competitively accurate in comparison with other algorithms. Moreover, via an evolutionary optimization process, the optimal number of features for each data set is determined. Through broad experimental evaluation, MMCC is proved to be significantly better compared to other well-known gene selection algorithms for 25 commonly used microarray data sets. Surprisingly, high accuracy in classification by Support Vector Machine (SVM) is achieved by less than10 genes selected by MMCC in all of the cases.

## 1. Introduction

It is beyond any shadow of doubt that cancer is a crucial disease that presents different challenges in both diagnosis and treatment [1]. Similar to most of diseases, cancer is also influenced by genes, thus, determination of genetic landscape and profile of this disease is of utmost importance and significance [2, 3]. Recent advancements in high-throughput technologies such as Next Generation Sequencing (NGS), Microarray, Mass Spectrometry (MS), and imaging assays and scans have paved the way for researches to identify genetic causes of diseases [4, 5, 6, 7]. These new high-throughput representations require proper computational methods and techniques in order to obtain knowledge from them precisely [8, 9, 10, 11, 2]. Although these new technologies are infinitely beneficial and advantageous, they might be suffering from an enormous problem named “Curse of Dimensionality” if they have high number of observed genes (dimensions) but low number of samples [12]. Fortunately, manifold feature selection methods have been proposed so far to deal with “Curse of Dimensionality” and subsequent over-fitting in learning process [12, 13]. Although we analyze the proposed method by microarray data sets, such an approach can be helpful in numerous other feature selection problems [14, 15, 16, 17]. In addition, feature selection generally is helpful to reduce the time and memory complexities which are always issues. Moreover, the increased complexity in time or memory is not only limited to time and space occupations, but it adversely influences the performance of the algorithms due to the noise effect and outliers [18]. Finally, feature selection is also important for visualization purposes.

Generally, feature selection methods can be categorized into two broad groups of classifier dependent (such as ‘wrapper’ and ‘embedded’ methods) and classifier independent (such as ‘filter’ methods). In filter methods feature selection and classification components are separated and selection of feature is based on some heuristic criteria and scoring. In contrast, wrapper and embedded methods take advantage of a classifier evaluation of a subset by training/test accuracy and determination of the structure of specific classes of learning models to enhance the feature selection process respectively [13].

Model et al. [19] utilized numerous feature selection methods for DNA methylation based cancer classification. Li et al. [20] applied feature selection for tissue classification based on gene expression. Zhang et al. [21] used Support Vector Machine (SVM) with non-convex penalty for gene selection in cancer classification. Similarly, Cawley et al. [22] proposed a sparse logistic regression with Bayesian regularization for the same challenge. Logsdon et al. [23] suggested gene expression network reconstruction by convex feature selection when incorporating genetic perturbations in order to discover novel network relationships. Fitzgerald et al. [24] proposed a second order dimensionality reduction method using minimum and maximum mutual information models in multidimensional neural feature selectivity. Piao et al. [25] utilized an ensemble correlation-based feature selection method for cancer classification with gene expression data. Another ensemble approach for feature selection proposed by Yassi and Moattar [26] is a robust and stable feature selection method for genetic data classification. Aguas et al. [27] has shown that how feature selection methods can be applied to achieve reliability in classification of viral sequences by host species, and to determine the vital minority of host-specific sites in pathogen genomic data. Manchon et al. [28] proposed novel features and a multiple classifier approach for identifying cancer associated mutations in the cancer kinome.

Interested readers can refer to six great surveys and reviews of feature selection in [17, 29, 30, 31, 32] and Jain et al. [33] and the references therein.

In this paper, we aim to use Maximum-Minimum Correntropy Criterion (MMCC) for filter-based feature selection. The main contributions are summarized as follow:

- Anew formulation based on MMCC is investigated for feature selection.
- The optimal parameters of MMCC are found by an optimizer.
- Along with MMCC parameters, the number of features to be selected is determined and we do not need to scrutinize the performance of MMCC by different desired number of features.
- Robustness, speed and stability of MMCC are explored by testing on numerous data sets.

## 2. Materials and Methods

### 2.1. Correntropy

In Information Theoretic Learning (ITL)[34], Correntropy is defined as a local similarity measure[35] and it has shown robustness in dealing with various type of noise and outliers [36]. Given two random vectors *x* ∈ ℝ*^N^* and *y* ∈ ℝ*^N^*, Correntropy is defined as

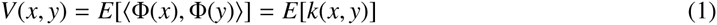

where *E*[.]is expectation operator, Ф(.) connotes a nonlinear mapping to high dimensional space, *k* is any kernel function and 〈.,.〉 denotes the inner product. The real world data sets usually contain a finite number of samples by which (1) can be restated as

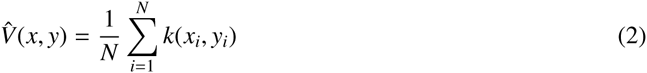

Kernel-based algorithms have shown great performance in the realm of machine learning. The kernel function utilized in Correntropy is often the Gaussian kernel defined as

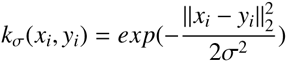

where *σ* is called the kernel width. In practice, *σ* can dramatically influence the performance of the algorithms based on Gaussian kernel. In this paper, the Gaussian kernel is utilized and its kernel width is determined by an optimizer which we will discuss in further sections.

The robustness of Correntropy has been investigated in face recognition [37], non-Gaussian signal processing [36], clustering [38] and classification [39], to name a few. To our knowledge, it is the first time that Correntropy is utilized for filter-based feature selection.

### 2.2. Filter-based Feature Selection via Maximum-Minimum Correntropy Criterion

Filter methods are predicated on a criterion J which measures the importance and usefulness of a feature or a subset of features. A potential criterion J would usually measure the correlation between the feature and the class label. Given a data set *X* and class label *Y*, we propose to use the *Maximum-Minimum Correntropy Criterion (MMCC)* score for a feature *X_k_*, i.e.

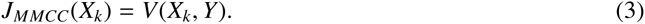

To select *K* features, one can rank features based on MMCC (3) and select the top *K* ones. However, the criterion in (3) suffers from selecting highly correlated features. Generally, the features are to be chosen that are individually *irrelevent* in order to avoid *redundancy* [13]. For this aim, the criterion in (3) is modified as

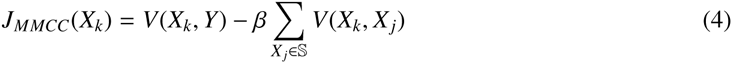

where *β* is a regularizer parameter and 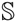 is the set of selected features. Note that we construct final selected features iteratively and in each iteration, one feature added to the selected set 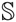. Parameter *β* is a trade-off between importance of similarity between feature *X_k_* and label and similarity between feature *X_k_* and already selected features.

There are three parameters (*β*, *σ* and number of features to be selected) which can influence the performance of the proposed problem. To find the optimal values for this parameters, an evolutionary process is utilized which we study in detail in further sections. Fig. 1 illustrates the procedure of MMCC that finds optimals values of three above-mentioned parameters and selects features.

**Figure 1:** The procedure of MMCC for selection optimal parameters and features.

**Figure 2:** Maximum test accuracy in 10,000 runs of SVM classification on selected features for 15 two-class data sets. The methods are MMCC, MIM [43], MRMR [44], CMIM [45] and JMI [46].

**Figure 3:** Maximum test accuracy in 10,000 runs of SVM classification on selected features for 10 multi-class data sets. The methods are MMCC, MIM [43], MRMR [44], CMIM [45] and JMI [46].

## 3. Results and Discussion

### 3.1. data sets

Throughout this article, in order to evaluate the proposed method we utilized 25 microarray data sets of gene expression levels adopted from [12]. 15 of these data sets are two-class and the rest 10 are multiclass. For each class, each sample represents a common phenotype or a subtype thereof. For convenience, these 25 data sets are assigned a reference number which the first 15 are for two-class and 16 to 25 are dedicated for multiclass data sets. The dimensions of these data sets vary from approximately 2000 to 25000 features and sample size varies approximately between 50 and 300 samples.

### 3.2. Performance

The optimal values of three parameters including *β*, Correntropy kernel width *σ* and the number of features to be selected can profoundly influence the performance of the proposed method. The classification accuracy is probably higher if we select more features. However, performing classification on data sets with more features requires more time and memory. On the other hand, if the number of selected features is not enought, the classification will fail, therefore, finding the number of selected features can be tricky. The performance of Correntropy is highly related to its kernle width. If the kernel width is low, Correntropy will operate similar to *l*_0_ norm and a part of data might be lost. In contrast, if the kernel width is high, Correntropy is approximately *l*_2_ norm which means that it loses its robustness in dealing with high corruption and non-Gaussian noise. The parameter *β* also adjusts the selected features’ redundancy.

To obtain these values, we utilize an adaptive version of Differential Evolution (DE) [40]. DE is one of the most powerful and effective population-based evolutionary computation methods for global optimization especially in continuous problems. To perform the optimization process, DE uses three operators including mutation, crossover and selection. Since these operators have their own parameters we applied jDE [41] which is an adaptive variant of DE. *jDE* requires a cost function for optimization tasks and one can use the accuracy of a classifier for this purpose. In this article, we utilize SVM with linear kernel available in Libsvm library [42].

Table 1 and Table 2 tabulate the optimal values of the above-mentioned parameters for two-and multi-class data sets, respectively. Further, we defined the lower and upper bounds of parameters of MMCC which have been brought in these tables.

**Table 1:** Optimal values of MMCC parameters for 15 two-class data sets.

**Table 2:** Optimal values of MMCC parameters for 10 multi-class data sets.

Next, the performance of MMCC is compared with 4 well-known filter methods: MIM [43], MRMR [44], CMIM [45] and JMI [46]. To make our comparison fair, obtained optimal number of features are set for each method. We divide each data set into train and test subsets: 80 percent of each data set dedicated to train classifier and the rest are for testing the performance of the SVM. In order to diminish the uncertainty of the SVM, we run it 10000 times independently. Tables 3 and 4 are stated the maximum accuracy of the SVM over these independent runs. For two-class data sets, in 4 of cases MMCC achieved the best performance among other methods and in 6 data sets, all methods obtained the same performance. In 3 of the data sets MMCC was the best with at least one of the other methods and MMCC had the poorest performance in 2 data sets.

**Table 3:** The time (in seconds) comparison of MMCC, MIM [43], MRMR [44], CMIM [45] and JMI [46] on 15 two-class data sets.

**Table 4:** The time (in seconds) comparison of MMCC, MIM [43], MRMR [44], CMIM [45] and JMI [46] on 10 multi-class data sets.

For multiclass data sets, in 3 of the cases MMCC had the best performance and in 2 data sets all methods performed equally. In 3 data sets, MMCC obtained the best results with at least one of the other methods and finally, in only one data set MMCC was not successful. It is important to note that some of the compared method evoked ‘out of memory’ error on big multiclass data sets. The list of selected genes are brought in Supplementary Material for interested readers.

### 3.3. Time

Filter-based feature selection methods (like mutual information based methods) usually needs to estimate several density functions. This estimation is usually depended to the number of samples and features and it becomes more times consuming as the dimension of data increase. Correntropy criterion, however, does not need to estimate any density functions and it is probably less time consuming. Experimental analysis on this issue illustrates that MMCC is significantly faster. Table (3) and (4) tabulate that time (in seconds) of the aforementioned methods. It is readily seen that the speed of the MMCC is competitive in comparison to other methods.

### 3.4. Stability

One of the most important aspects of any feature selection method is its stability. Following [47], stability of a feature selection method means the robustness of the feature preferences it suggests to differences in diverse training sets obtained from the same generating distribution. In fact, for a given data set, the stability of a method is the stability of the appearance of certain features after resampling. Let *Y* be set of all features and |*Y*| be the cardinality of it, according to Dunne et al. [48], Average Normalized Hamming Distance (ANHD) is a stability measure for a feature selection method as follow:

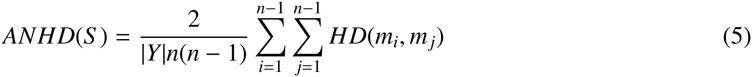

Here, HD is the hamming distance of mi and mj which are two binary vectors corresponding to two subsets Si and Sj from two different samples of a data set originated from n runs, i.e.

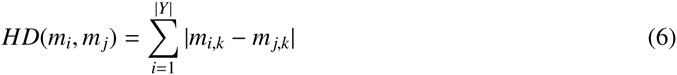

Where,

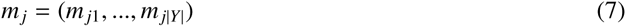

In this section, we utilized ANHD in order to evaluate the stability of MMCC over *n* = 10 independent runs and in each run 90% percent of the samples were considered randomly for evaluation of MMCC. The results are stated in Table (5). As illustrated in these tables, in almost all of the data sets MMCC reached an ANHD almost near to 0 which indicates minimum variation. Therefore, MMCC is not only fast and robust against noise and outliers but also significantly stable which is vital and advantageous for a feature selection algorithm.

**Table 5:** Exploring the stability of 15 two-class (above) and 10 multi-class (bottom) data sets via Average Normalized Hamming Distance (ANHD) [47].

## 4. Conclusion

In this paper, we proposed Maximum-Minimum Correntropy Criterion (MMCC) for selection of informative genes from microarray data sets. We have shown that by less than 10 genes among thousands ones it is possible to achieve significantly high accuracy in cancer classification. Furthermore, we stated numerous advantageous for MMCC such as low computational cost, robustness (because of correntropy) and more importantly its stability (evaluated based on Average Normalized Hamming Distance (ANHD)). Our experiments show that, genes selected by MMCC achieved maximum performance in cancer classification by SVM in almost all of the data sets in comparison with other well-known compared feature selection algorithms. Interestingly, some of these algorithms faced “out of memory” error during gene selection process especially for multiclass data sets, however, MMCC overcomes this problem as well which indicates that it is required less memory compared to other algorithms.

